# High-resolution quantification of the rhizosphere effect along a soil-to-root gradient shows selection-driven convergence of rhizosphere microbiomes

**DOI:** 10.1101/2024.06.21.600027

**Authors:** Sanne W.M. Poppeliers, Juan José Sánchez-Gil, José Luis López, Bas E. Dutilh, Corné M. J. Pieterse, Ronnie de Jonge

**Affiliations:** Plant-Microbe Interactions, Department of Biology, Science for Life, Utrecht University, 3584CH, Utrecht, The Netherlands; Theoretical Biology and Bioinformatics, Department of Biology, Science for Life, Utrecht University, 3584CH, Utrecht, The Netherlands; Instituto Andino Patagónico de Tecnologías Biológicas y Geoambientales, Bariloche, Rio Negro, Argentina; Institute of Biodiversity, Faculty of Biological Sciences, Cluster of Excellence Balance of the Microverse, Friedrich-Schiller-University Jena, 07743, Germany; AI Technology for Life, Department of Computing and Information Sciences, Department of Biology, Utrecht University

**Keywords:** Plant root microbiome, rhizosphere effect, soil-root gradient, priority effects, natural selection

## Abstract

Plants secrete a complex array of organic compounds, constituting about a third of their photosynthetic products, into the surrounding soil. As a result, concentration gradients are established from the roots into the bulk soil, known as the rhizosphere. Soil microbes benefit from these root exudates for their survival and propagation, and consequently, the composition of the rhizosphere microbial community follows the gradient of available compounds, a phenomenon oftentimes referred to as the rhizosphere effect. However, the fine-grained changes in the microbial community along this soil-root gradient have not been well described. Yet such insights would enable us to underpin the ecological rules underlying root microbial community assembly. Therefore, here we harvested the roots of individual *Arabidopsis thaliana* plants grown in three different natural soils at high-resolution, such that we could interrogate community assembly and predict microbial growth rate across consecutive, fine-grained, rhizosphere ‘compartments’. We found that the strength of the rhizosphere effect depends on root proximity and that microbial communities closer to the roots harbour related microbes. Closer to the roots, microbial community assembly became less random and more driven by selection-based processes. Intriguingly, we observed priority effects, where related microbes that arrive first are more likely to establish, and that microbes might use different ecological growth strategies to colonise the rhizosphere. All effects appeared to be independent from starting conditions as microbial community composition converged on the root despite different soil ‘microbial seed banks’. Together, our results provide a high-resolution view of the microbiome changes across the soil-root gradient.

## Introduction

As molecules diffuse through a matrix, they dilute as they move further away from the source. The same principle holds for plant roots and their exudates. Plants secrete a complex array of organic compounds into the soil^1^, about a third of their photosynthetic products^2^. As a result, concentration gradients are established from the roots into the bulk soil and vice versa^3^. For example, in ryegrass, the concentration of soluble organic carbon compounds including soluble sugars and oxalic acid decreased further away from the roots^4^. The soil surrounding the roots that is influenced by root activity is called the rhizosphere^5^. Its shape and size depend on plant properties and soil type^6^. The rhizosphere is both dynamic and gradual, as roots continue to grow and explore new parts of the surrounding soil and microhabitat properties gradually change from root to bulk soil^3^.

Because the rhizosphere is a carbon-rich environment, it is a hotspot for microbial activity^7,8^. Rhizosphere microbial communities are often different from the surrounding bulk soil, a pattern that is referred to as the ‘rhizosphere effect’^9–13^. The root-associated microbiome has a higher cell density^15,16^, is often less diverse, and harbours a distinct set of species^14,15^. Since soil microbes are (partly) dependent on root exudates for their survival, rhizosphere microbial community composition most likely follows the gradient of available metabolites, gasses, ions, and other substances^3,16^. Many, diverse rhizosphere-competent taxa that contribute to the separation between bulk soil and rhizosphere microbial communities have been identified^17^, yet how microbial community composition differs on smaller scales along the soil-root gradient is still largely unknown.

The rhizosphere effect in *Arabidopsis thaliana* (hereafter Arabidopsis), an extensively studied model plant, is often found to be relatively small^18–21^, especially compared to plant species with a longer life history^22^ or crops like wheat, oat, and pea^21^. In these studies, the rhizosphere is defined as ‘loosely adhering soil’^23^. However, evidence points out that for microbial communities firmly attached to the root (often referred to as the ‘rhizoplane’), the rhizosphere effect in Arabidopsis is comparably strong^22^. It is likely that the observed rhizosphere effect not only depends on the amount and composition of root exudates and their diffusion rates through the soil, but also on how the rhizosphere is collected. For Arabidopsis, which has relatively small roots compared to large crops such as maize and wheat, different sampling strategies might be necessary to reveal the scope of the root influence, in order to fully explore how rhizosphere microbial communities are assembled, and assess what taxa and functions are associated with rhizosphere competence.

To better understand establishment of the rhizosphere microbial community, it is essential to characterize processes underlying its assembly. Traditionally, microbial community assembly is most often explained by mechanisms of selection^24^, while more and more studies show that in complex communities, chance, and environmental heterogeneity can be equally important^25–28^, also in the rhizosphere^29,30^. This might be especially relevant when trying to engineer plant microbiomes using plant-beneficial microbes^31^. Given that the root microbiome is mainly formed by a variety of microbes present in the surrounding bulk soil^32^, as well as those originating from seeds or carried by the air^33–35^, it is probable that random processes have a greater impact in the rhizosphere than, for instance, the mammalian gut, where numerous microbes are transmitted during birth^36^. Who ends up in the rhizosphere is primarily driven by the resident soil microbial community, and who can colonize first^37^, and only secondarily through selection by the plant, depending on root proximity. While there is a substantial amount of research on taxa, genes or traits that are enriched in the rhizosphere under specific circumstances, it is still not clear what ecological community assembly processes steer this environment^38^.

With this study we wanted to determine the strength and dynamics of the rhizosphere effect in Arabidopsis across the soil-root continuum, characterize the changes in microbial community composition along this gradient and assess the importance of selective and random community assembly processes. To do so, we grew Arabidopsis plants in natural soil sampled from the same geographic location over three different years and harvested the rhizosphere in a reproducible manner such that the microbiome in each consecutive ‘compartment’ could be measured and compared to the microbes in the unplanted bulk soil. We focused on the bacterial portion of the rhizosphere microbiome, the largest microbial portion^39^, and found that the strength of the rhizosphere effect in Arabidopsis is dependent on root proximity. Bacterial communities closer to the roots harbour phylogenetically similar microbes, even though different microbes are detected across samples, a phenomenon that we attribute to the influence of priority effects. As we sampled closer to the roots, we found that microbial community assembly became less random and was more driven by selection-based processes, and this effect was largely independent from starting conditions as microbial communities converged on the root despite different bulk soil ‘microbial seed banks’.

## Materials and Methods

### Soil collection and preservation

The soil used in this study was taken from the Reijerscamp nature reserve, the Netherlands (52°01′02.55″, 5°77′99.83″), where previously an abundant endemic Arabidopsis population was found^73^. Agricultural practices ended in 2000, and subsequently the area was restored as natural grassland. The soil was described by Berendsen et al. 2018^73^ as a gleyic placic podzol, consisting of coarse sand and gravel covered by a 30–50 cm top layer. The top 20 cm of soil was collected in October 2018 for experiment 1 (hereafter exp 1), April 2019 for experiment 2 (exp 2) and December 2019 for experiment 3 (exp 3), air dried and sieved (1 cm sieve) to remove plant debris and rocks, and subsequently stored at room temperature. Prior to the experiment the soil was mixed with 100 mL of tap water per kg of dried soil.

### Experimental set-up

*Arabidopsis thaliana* accession Col-0 seeds (N1093; Nottingham Arabidopsis Stock Centre) were surface sterilized by fumigation using a mixture of 100 mL bleach and 3.2 mL 37% HCl for 4 h. Seeds were sown on 1% Murashige and Skoog (MS) medium^74^ with 0.5% sucrose, and after 2 days of stratification in the dark at 4°C, the seeds were transferred to a growth chamber (21 °C, 70% relative humidity, 10 h light/14 h dark, light intensity 100 μmol · m^−2^ · s^−1^) to allow germination. Two-week-old seedlings were transferred to individual 60-mL pots with approximately 100 g of Reijerscamp soil. Bulk soil pots were left unplanted. Pots were watered for 3 weeks every other day and during the whole experiment developing seedlings from plant seeds that came with the soil were removed with tweezers upon detection.

Five-week-old Arabidopsis roots were harvested using a standardized sampling method (**Supplementary Figure 1a**), resulting in four compartments of the soil-root gradient, in which in each consecutive compartment microbiome samples were harvested in increasingly closer proximity to the root (**Supplementary Figure 1b**). Exp1 was a pilot where each sample consisted of two plants, while in exp2 and exp3 we used one plant per sample, and more replicates were added to accommodate in-depth analyses. For rhizosphere-4 samples (RS4), roots were taken out of the soil and shaken briefly until most loose soil fell off and stored in a 2-mL Eppendorf tube. For rhizosphere-3 samples (RS3) this was done similarly, and additionally roots were stripped of as much soil as possible with clean gloves and by tapping the roots on clean paper before transferring them to a 2-mL Eppendorf tube. For rhizoplane-2 samples (RP2), roots that were harvested the same as for the RS3 samples, were transferred to a Falcon tube containing 25 mL MgSO_4_ and inverted 5 times by hand. Roots were then taken out, carefully dried with paper, and put in a clean 2-mL Eppendorf tube. For rhizoplane-1 samples (RP1), roots were harvested according to the RS3 samples, and transferred to a Falcon tube containing 25 mL phosphate-Silwet (PBS-S) buffer (per L 6.33 g NaH_2_PO_4_.H_2_O, 10.96 g Na_2_HPO_4_.2H_2_O and 200 µL Silwet L-77) and vortexed on maximum speed for 15 s. After this, the roots were transferred to a new Falcon tube with 25 mL PBS-s buffer and vortexed again for 15 s. Roots were than taken out, dried with paper, and transferred to a clean 2-mL Eppendorf tube. Up to 0.25 g of bulk soil was taken from each unplanted pot. Root or soil samples were flash frozen in liquid N_2_ directly after sampling and stored at -80°C.

### DNA isolation and 16S amplicon sequencing

DNA extractions for exp1 and exp2 were done with the Mo Bio PowerSoil kit (Qiagen, Germantown, USA) according to the manufacturer’s instructions with one adjustment optimized for Reijerscamp soil: after addition of lysis buffer and solution C1, samples were additionally incubated at 70°C for 10 min. The remaining steps were performed as described by the manufacturer’s instructions. For exp3, DNA extractions were performed with the MagAttract PowerSoil DNA kit (Qiagen, Cat. No. 27000-4-KF) optimized for the KingFisher™ Flex Purification System (Thermo Scientific, 183 Waltham, MA, USA). DNA quality and quantity was checked using a NanoDrop 1000 spectrophotometer (Thermo Scientific, 183 Waltham, MA, USA). For exp 1, we pooled DNA of two individual samples from the same compartment. All sampled were normalized to 5 ng · µL^-1^. Subsequently, we amplified the hypervariable V3-V4 region of the 16S rRNA gene with phasing primers CCTACGGGNGGCWGCAG and GACTACHVGGGTATCTAATCC (**Supplementary Table 3**), using the standard protocol from Illumina. In short, the amplification reaction mixture contained 2.5 µL of sample, 5 µL of primer mix, 5 µL PCR grade water and 12.5 µL KAPA HiFi HotStart ReadyMix (Roche, Indianapolis, USA). The PCR conditions were 98°C for 5 min, 25 cycles of 98°C for 40 s, 55°C for 30 s, 72°C for 60 s and a final extension at 72°C for 10 min. For the index PCR the reaction mixture contained 2.5 µL of DNA from PCR 1,

2.5 µL of each primer, 5 µL PCR grade water and 12.5 µL KAPA HiFi HotStart ReadyMix (Roche, Indianapolis, USA). Samples were normalized and pooled in equimolar amounts and the pools were sequenced on the Illumina MiSeq machine with the 2 x 300 bp V3 kit at the USEQ sequencing facility (Utrecht University, the Netherlands).

### Bioinformatics and statistical analysis

Raw sequencing data was processed with Qiime2 version 2019.7^75^. Each individual sequencing run (1 for exp 1, 2 for exp 2 and exp 3) was processed separately as follows. After importing the data, primers were removed using *Cutadapt* 2.8^76^. Raw sequence data were demultiplexed and quality filtered using the q2-demux plugin followed by denoising with DADA2^77^ (via q2-dada2) where sequences were clustered into Amplicon Sequence Variants (ASVs) at >99% identity. Singletons and doubletons were removed, and data from individual sequencing runs was merged. Taxonomy was assigned to the ASVs based on the SILVA reference database (99% similarity version 132) using the classify-consensus-vsearch plugin (VSEARCH 2022.8.0)^78,79^. After pre-processing of the sequencing data, ∼2.23 · 10^7^ reads remained and were clustered into 35762 ASVs from three experiments and 225 samples. Mean number of reads per sample was 99,092 reads, and 8 samples per compartment for exp1, and 18 samples per compartment for exp2 and exp3 were retained. All subsequent analyses were executed in R Statistical Software (v4.1.3; R Core Team 2022). The package *Phyloseq*^80^ was used to obtain the number of plant reads, read depth, and α- and β-diversity, where plant reads (chloroplasts and mitochondria) were removed before analysis. From the top 50 ASVs that most strongly influence the ordination, we calculated their association with PC1 as abs(PC2). The Sloan neutral model for prokaryotes^25^ was implemented on a rarefied feature table (10,000 reads per sample), using an adjusted publicly available script by Burns and co-authors (2016)^26^. For estimation of abundance-weighted growth rate potential we largely followed Lopez et al. 2023^37^. In short, we first extracted genus-level estimates from the EGGO database^81^ which includes predicted minimum doubling times (PMDTs) for over 217,000 prokaryotic genomes and computed the median PMDT for each genus. Next, we summed the abundances of each ASV within a genus, and computed abundance-weighted growth rate potential for each sample type by multiplying the relative abundance of each genus with the genus-specific PMDT and taking the median of those values as representative for a specific compartment.

### Data availability

The raw 16S rRNA amplicon sequencing data generated and used in this study have been deposited at the NCBI Sequence Read Archive (BioProject: PRJNA1117364) and are publicly available as of the date of publication.

## Results

### Similar plant phenotypes and starting microbiomes with different soil batches

To study the rhizosphere effect in Arabidopsis along a soil-root gradient, we conducted three experiments in which we sampled roots in similar ways. We assessed the reproducibility of the rhizosphere assembly process by using three comparable soil batches obtained from the same natural site over multiple years (the Reijerscamp nature reserve, 52°01′02.55″, 5°77′99.83″; **Figure 1a**). Soil batches were harvested in different seasons: for experiment 1 (exp1) and experiment 3 (exp3) material was collected in the fall (October 2018 and December 2019 respectively) and for experiment 2 (exp2) this was during spring (April 2019). To evaluate whether the different soil batches affect plant performance, - this could influence our ability to assess rhizosphere microbiome assembly reproducibility -, we measured plant biomass in each experiment. We note that plants grown on all batches displayed similar growth with only limited variation between experiments. Shoot weight was ∼1.5 times higher in exp2 than in exp1 and exp3 which had similar shoot weights (**Figure 1b**; ANOVA, p<0.05).

**Figure 1.**
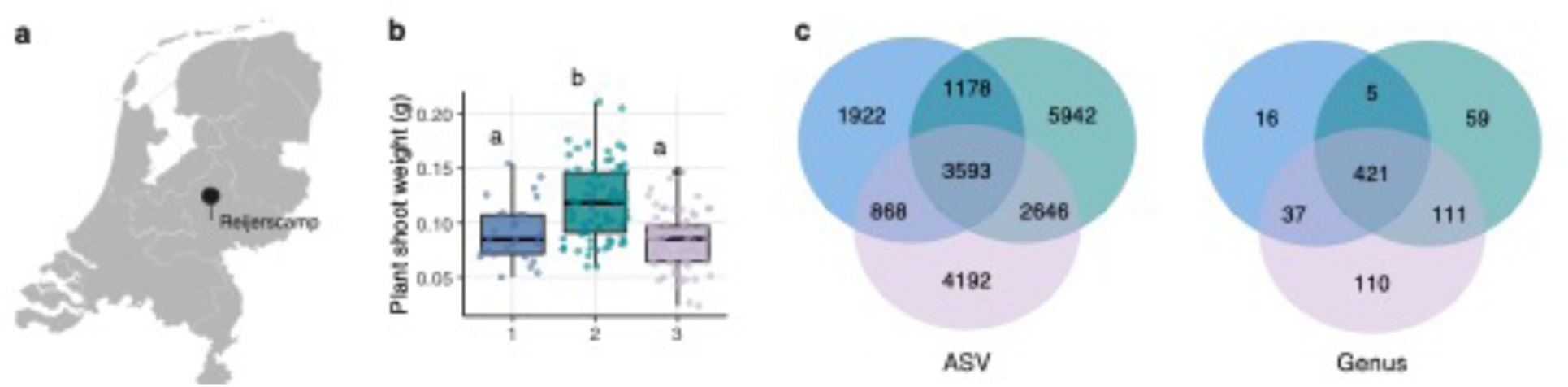
Similar plant phenotypes and starting microbiomes between different soil batches. **a** Sampling location of the Reijerscamp nature reserve in the Netherlands. **b** Plant shoot weight per experiment. Plants from exp2 had ∼1.5x higher shoot weight than plants from exp1 and exp3. **c** Overlap in number of ASVs (left) and genera (right) per experiment. For ASVs, 40% was found in ≥ 2 experiments, while 75% of all genera in our dataset occurred in ≥ 2 experiments.

For each experiment, we systematically collected the root-soil interphase in five high-resolution ‘compartments’ along the soil-root gradient: soil (S), rhizosphere-4 (RS4), rhizosphere-3 (RS3), rhizoplane-2 (RP2) and rhizoplane-1 (RP1) (**Supplementary Figure 1**, see Materials and Methods), and profiled the bacterial communities by amplicon-based sequencing targeting the V3-V4 region of the *16S rRNA* gene. To evaluate whether the different soil batches have comparable microbial communities, we first evaluated the overlap in amplicon-sequence variants (ASVs) and genera in these soils. Of all ASVs, 40% was found in ≥ 2 experiments, while 75% of all genera in our dataset occurred in ≥ 2 experiments (**Figure 1c**). The ASVs identified in the soil samples of the different experiments were mostly unique, but phylogenetically similar due to a high overlap at the genus level, thus indicating comparable microbial starting conditions across all three experiments.

### Strength of rhizosphere effect gradually increases towards the root

With comparable microbial starting conditions established, our focus shifted to a more detailed examination of the soil-root continuum. Moving progressively from RS4-RP1, we sampled roots with less adhering soil as sample weight decreased significantly between consecutive compartments for exp2 and exp3 and was on average 4.7±0.9 times lower in RP1 compared to S (Figure 2a). Consequently, the microbial community was expected to be increasingly influenced by the plant when sampling from RS4 to RS1. Because these consecutive compartments include the plant root itself (**Supplementary Figure 1**, see Materials and Methods), we could use the amount of plant reads in our samples as a proxy for root distance. The percentage of plant reads (assigned taxonomy either ‘chloroplast’ or ‘mitochondria’) in our samples gradually increased when sampling closer to the roots (S<RS4<RS3<RP2<RP1; 0.157±0.215% in S to 37.0±14.4% in RP1) providing a data-driven way to quantify the distance to the roots (Figure 2b, **Supplementary Table 1**; Kruskal-Wallis, p<0.001).

**Figure 2.**
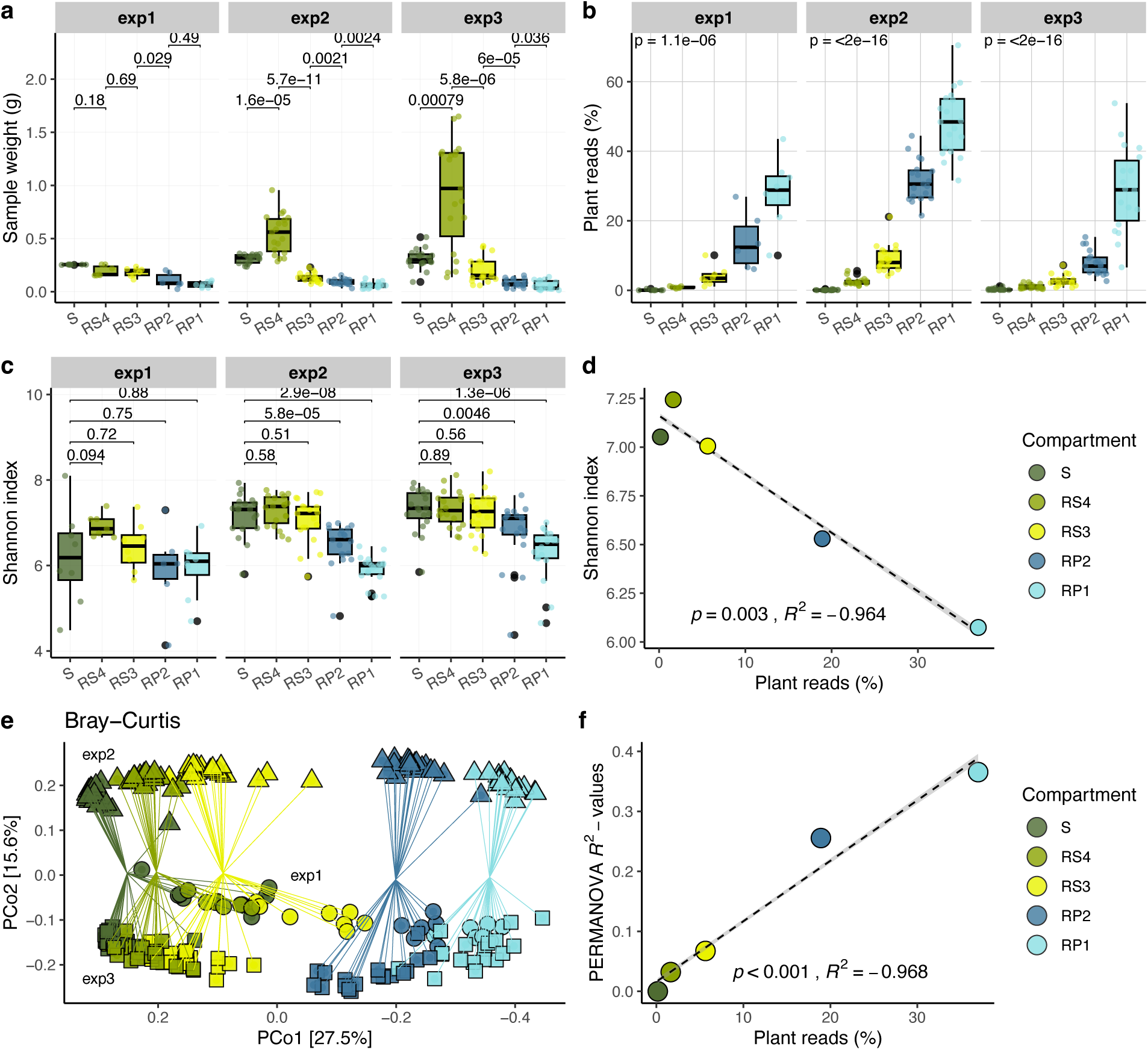
Stronger rhizosphere effect closer to the root. **a** Sample weight in grams for each compartment in each experiment. Numbers indicate p-values with Bonferroni-correction for multiple testing (Wilcoxon test). **b** Percentage of plant reads in each compartment (Kruskal-Wallis, *p* < 0.001, **Supplementary Table 1**). **c** Shannon index for each compartment in each experiment depicting the microbial diversity per sample. This index integrates both species evenness and species richness. **d** Strength of the rhizosphere effect quantified by the reduction in microbial diversity (Shannon index) closer to the root. The percentage of plant reads is used as a proxy for sample-root distance. **e** Principal coordinate analysis (PCoA) plot based on Bray-Curtis dissimilarity highlighting reproducibility of the soil-root gradient sampling (PCo1) across multiple experiments (PCo2). **f** The pairwise PERMANOVA *R*^2^-values of the compartment community composition difference compared to soil are used as an indicator of the strength of the rhizosphere effect (Pearson; *p* < 0.001, *R*^2^ = 0.984). Compartments are indicated by abbreviations: soil (S), rhizosphere-4 (RS4), rhizosphere-3 (RS3), rhizoplane-2 (RP2) and rhizoplane-1 (RP1).

Using the microbiome datasets sampled along the soil-root gradient, we then tested whether the alpha diversity (hereafter α-diversity) within our microbial communities decreased from bulk soil to rhizosphere and rhizoplane by calculating and comparing the Shannon index. The Shannon index is mostly driven by evenness^40^ and is more robust to inherently noisy *16S rRNA* ASV data^41^. In exp2 and exp3, the α-diversity decreased relative to S for the rhizoplane (RP2 and RP1), with the lowest values for the RP1 samples (Figure 2c; Wilcoxon test, *p* < 0.05). α-diversity, which can be seen as a measure for the strength of the rhizosphere effect, correlates linearly with the percentage of plant reads, i.e. with the distance to the roots (Figure 2d; Spearman, *p* = 0.003, *R*^2^ = -0.982).

To compare the microbial community composition along the soil-root gradient between experiments, we measured between-sample beta-diversity (hereafter β-diversity) between all pairs of samples using the Bray-Curtis dissimilarity measure and used these as input for a Principal Coordinate Analysis (PCoA) to visualize the differences. We then plotted the samples from all five compartments along the first two principal coordinates (PCos) of the PCoA. The distance to the plant root was the main factor that separated the samples along PCo1, which explained 27.5% of the variation in the dataset (Figure 2e). Differences between the three experiments with different soil batches represented the second most important factor along PCo2 (15.6% explained variation), and exp1 and exp3 were more similar to each other than exp2. Exp1 and exp3 were performed with soil batches collected from the same season (fall), while the soil from exp2 was collected in spring which might explain these observations. The results of pairwise permutational multivariate analysis of variance (PERMANOVA) using 999 permutations showed that there is a clear difference in the microbial community composition for each compartment-experiment combination (Adonis, *p* = 0.001). Although the rhizosphere effect is significant for RS4 samples which are most similar to S as they contain most soil, the effect size is small (*R*^2^ = 0.027). The microbiome of the RS3, RP2 and RP1 samples progressively differed from S (*R*^2^ = 0.063, 0.26 and 0.37). Similar as for α-diversity, the *R*^2^-values that represent the strength of the rhizosphere effect correlate very well with the percentage of plant reads (Pearson, *p* < 0.001, *R*^2^ = 0.984; Figure 2f). In conclusion, our sampling strategy effectively captures various high-resolution zones along the soil-root gradient and reinforces that the rhizosphere effect, wherein root activity influences the microbiome, strengthens gradually towards the root.

### Proteobacteria and Bacteroidetes replace other phyla on the roots

Our results show that the community composition of the compartments increasingly changes along the soil-root gradient, and that rhizoplane (RP2 and RP1) compartments have significantly lower diversity than soil samples. To explore what taxa are mainly causing this decrease in diversity we calculated the relative abundance of the major phyla that were present in all experiments (more than 5% relative abundance in a compartment within an experiment, Figure 3Error! Reference source not found.**a**). In line with previous studies^18,19,22^, *Proteobacteria* were on average the most abundant phylum across all experiments (39.0±11.2%) and their relative abundance was 1.9 times higher in RP1 compared to S (Wilcoxon test, *p* < 0.001). This was mainly due to expansion of the families *Burkholderiaceae* and *Oxalobacteraceae*, which together made up ∼21.0±4.5% and ∼15.2±7.3% of all Proteobacteria in RP1, respectively (**Supplementary Figure 2**). Although *Bacteroidetes* were not the most abundant phylum (9.96±3.95%), they showed the largest increase in relative abundance: 2.3 times higher in RP1 relative to S (Figure 3b; Wilcoxon test, *p* < 0.001). The largest increase on the root was due to the family of *Sphingobacteriaceae*, which became 9.3 times more abundant in RP1 compared to S.

**Figure 3.**
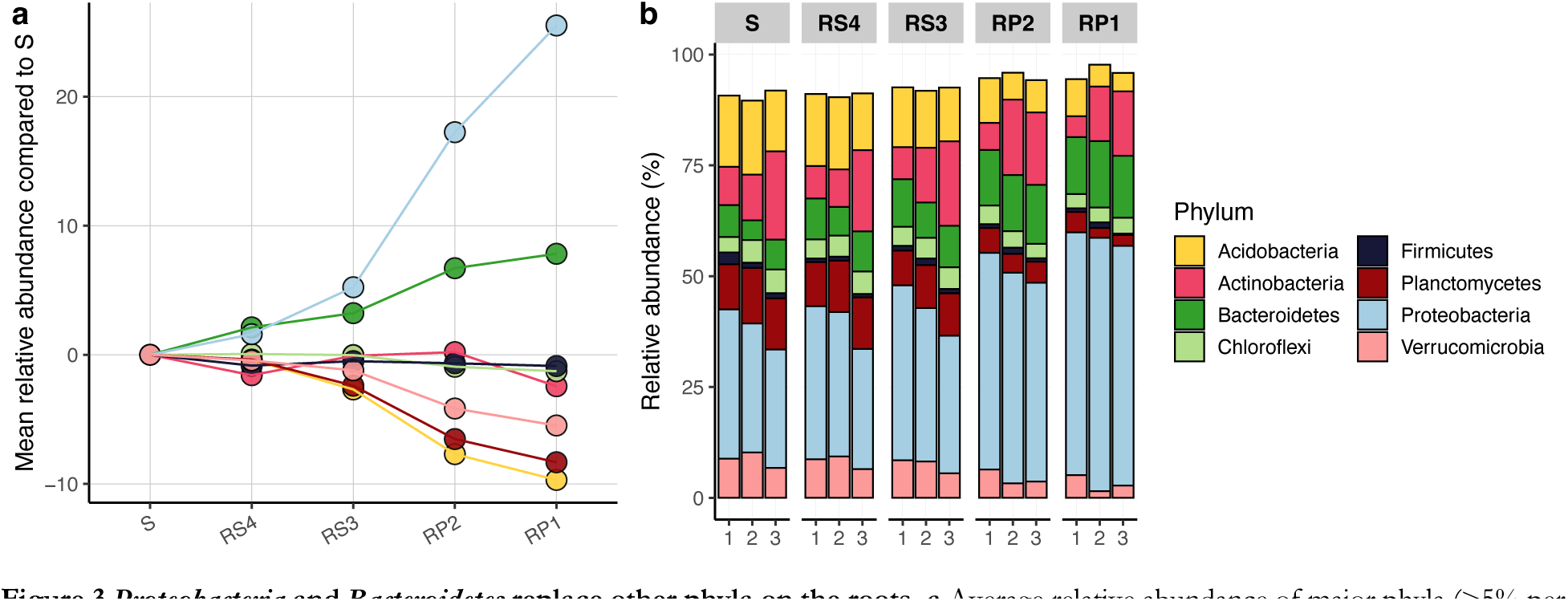
*Proteobacteria* and *Bacteroidetes* replace other phyla on the roots. **a** Average relative abundance of major phyla (>5% per compartment per experiment) minus their abundance in S in each soil-root compartment. **b** Relative abundance of major phyla. Remaining percentage of abundance is occupied by other phyla. Numbers on the x-axis indicate different experiments. Compartment abbreviations: soil (S), rhizosphere-4 (RS4), rhizosphere-3 (RS3), rhizoplane-2 (RP2) and rhizoplane-1 (RP1).

Several taxa decreased in relative abundance towards the root, including *Acidobacteria*, *Planctomycetes* and *Verrucomicrobia*. These phyla had comparable abundances in soil (15.4±2.1%, 11.7±2.1% and 8.6±2.0%, respectively) which decreased towards the root in a similar way, resulting in their abundance being ∼3 times lower in ‘rhizoplane’ samples (5.2±2.2%, 2.7±1.7% and 2.7±1.4%, respectively; Wilcoxon test, *p* < 0.001). Possibly, *Acidobacteria*, *Planctomycetes*, and *Verrucomicrobia* exhibit similar life strategies that are adapted to soil specifically and therefore they could be particularly susceptible to competition from rhizosphere-competent taxa. The relative abundance of *Actinobacteria*, *Firmicutes* and *Chloroflexi* varied less between compartments and stayed relatively constant along the soil-root gradient. Thus, it appears that *Proteobacteria* and *Bacteroidetes* replace, specifically, *Acidobacteria*, *Planctomycetes* and *Verrucomicrobia* closer to the roots.

To better understand which specific taxa from the *Proteobacteria* and *Bacteroidetes* increase in abundance on the plant root, we quantified the contribution of each ASV to the first two PCos in Figure 2e. We visualized this in a biplot that displays the top 25 ASVs that are most influential in differentiating the samples (**Supplementary** Figure 3), and quantified which ASVs correlate most strongly with PCo1 (**Supplementary Table 2**). We found that while *Streptomyces* and *Sphingomonas* ASVs most strongly influence the ordination, the abundance of the proteobacterial genera *Massilia* and *Devosia* is strongly associated with PCo1, along which the samples from the different compartments separated. *Massilia* constitutes a significant proportion of all *Proteobacteria* (∼12.9±6.9%) and was the most abundant genus in the RP1 communities (∼7.4±4.4%). This indicates that *Massilia* and *Devosia* are most enriched close to the roots and can be considered rhizosphere competent, i.e., capable of surviving and thriving in the rhizosphere.

### Rhizosphere microbiomes converge from diverse starting points

We observed that rhizosphere microbiomes become less diverse and more divergent from the soil microbiome as we move closer to the plant root (Figure 2), and that select taxa are repeatedly enriched along this soil-root gradient in different experiments (Figure 3; **Supplementary Figure 3**). Together, these results suggest that the enrichment of taxa towards the plant root is non-random and related to the genetic and functional similarity of the enriched taxa. To test this hypothesis, we calculated between-sample differences in taxonomic composition using the weighted UniFrac metric. As opposed to the previously used Bray-Curtis dissimilarity metric, which considers ASVs as categorical units, weighted UniFrac additionally incorporates the phylogenetic distances between ASVs, thereby accounting for their overall relatedness. When we then perform PCoA on these between-sample differences, we observed that samples from different experiments converged towards the roots (Figure 4a). Thus, the rhizosphere-associated microbes were phylogenetically similar in independent experiments, reflecting their genetic and functional similarity^42^. This is further supported by the decrease of Faith’s phylogenetic diversity^43^ (determined by summed branch lengths of the 16S rRNA V3-V4 amplicon-based phylogenetic tree) in the RP2 and RP1 samples, showing that communities near the roots contain phylogenetically more similar species than those in soil (Wilcoxon test, *p* > 0.05; Figure 4b). This suggests that the Arabidopsis rhizosphere is a selective environment consistently populated by a phylogenetically and likely functionally similar, competitive pool of microbes.

**Figure 4.**
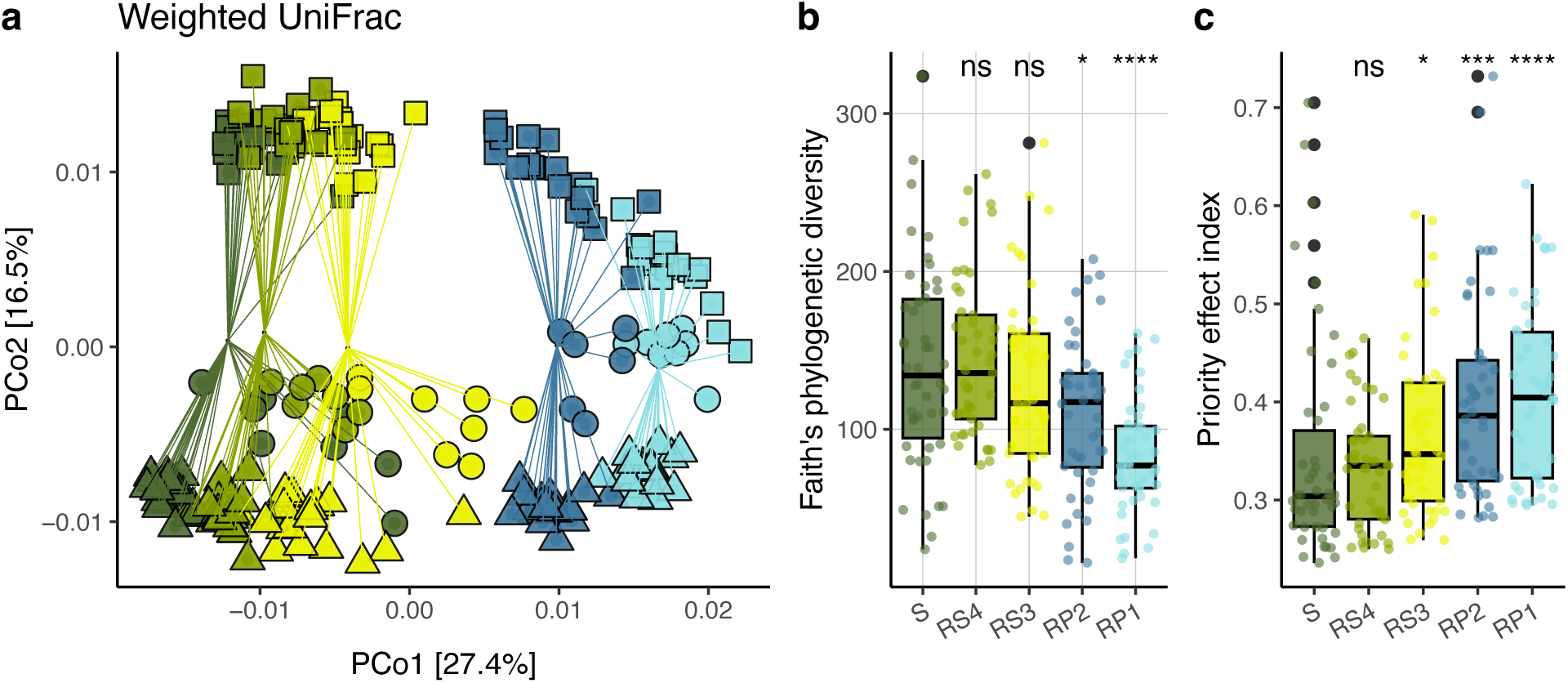
Phylogenetic clustering of micro-organisms closer to the roots which appears to be driven by priority effects. **a** PCoA based on weighted UniFrac, which incorporates the phylogenetic distances between ASVs, thereby accounting for their relatedness. **b** Faith’s phylogenetic diversity (determined by summed branch lengths of the 16S rRNA V3-V4 amplicon-based phylogenetic tree) per compartment. **c** Euclidian distances for all samples to the compartment centroid with the Bray-Curtis dissimilarity measure. A greater distance to the centroid implies greater dissimilarity between communities, indicating a higher number of distinct ASVs that are present between those samples. **d** Euclidian distances for all samples to the compartment centroid with the weighted UniFrac dissimilarity measure, where phylogenetic relatedness of ASVs is considered. For all panels the Wilcoxon test was performed. * *p* < 0.05 ** *p* < 0.01 and **** *p* < 0.0001, ns = not significant. Compartment abbreviations: soil (S), rhizosphere-4 (RS4), rhizosphere-3 (RS3), rhizoplane-2 (RP2) and rhizoplane-1 (RP1).

To further confirm the presence of a taxonomic signal across the soil-root gradient we quantified the degree of sample clustering by compartment that can be visually observed in the PCoA plots (Figure 3e, Figure 4a). To this end, we calculated the average Euclidian distance to the compartment centroid for each compartment using both Bray-Curtis and weighted UniFrac metrics and combined those metrics into a so-called ‘priority effect index’ by subtracting the weighted UniFrac values from those calculated using Bray-Curtis. A larger distance to the compartment centroid indicates more variation between samples of the same compartment and suggests that priority effects have played a larger role closer to the root. We found that the distance to the compartment centroid increased slightly in the RS3 compartment, and more in the rhizoplane samples relative to S (Figure 4c; Wilcoxon test, *p* < 0.05). We hypothesize that priority effects play a role in root colonization, especially in the rhizoplane, as phylogenetically related taxa that by chance arrive on the root first might have a competitive advantage. This would cause samples to diverge on the individual or ASV level as these are functionally redundant but converge on higher taxonomic ranks (Figure 4a).

### Rhizosphere effect is associated with niche-based community assembly

Our results suggest that microbial communities near plant roots experience selective pressure, resulting in phylogenetically similar communities. Thus far, the contribution of selection to the shaping of microbial communities along the soil-root gradient has not been quantified. Four main ecological processes govern microbial community assembly: selection, dispersal, drift, and speciation^44^. Traditionally, biodiversity is explained by niche theory, which proposes that each species has unique traits that allow it to occupy a specific environment^45^. This theory assumes that species are fundamentally different and can coexist because of these differences. In contrast, neutral theory explains diversity as a balance between random dispersal, drift, and/or speciation^46,47^. Both niche and neutral processes can occur simultaneously in the assembly of local communities^48^. To investigate to what extent these processes contribute to rhizosphere microbiome assembly along the soil-root gradient, we assessed the fit of the *Sloan Neutral Community Model for Prokaryotes* to the distributions of ASVs in our soil-root gradient data^25^. In short, the model predicts that, under a neutral sampling process, taxa that are abundant in the metacommunity are also widespread in local communities, while rare taxa are likely to be lost due to ecological drift^26^. Here, for each compartment, we compare the observed prevalence of ASVs (percentage of samples in which an ASV occurs) in the local root-compartments to their relative abundance in the summed bulk soil samples, using a β-distribution where the parameter *m* is fitted.

The model fit varied between compartments, with a considerably lower fit for the washed ‘rhizoplane’ roots (RP2 and RP1) compared to the ‘rhizosphere’ compartments (RS4 and RS3; Figure 5a). A lower model fit indicates that community composition is primarily non-neutral, i.e., driven by selection, or non-random dispersal or speciation, although the latter process is likely not relevant in our 21-day experiment. The model fit was almost zero for RP1 samples, indicating that the microbial communities in this compartment are strongly selected for or dispersal enriched. The estimated migration rate *m* decreased towards the root, suggesting that microbial communities near the roots are not shaped by dispersal from the bulk soil, but rather by reproduction and replacement from within the local community (Figure 5b). The model fit was positively correlated with the percentage of plant reads (Figure 5c). In all compartments, there were ASVs that occurred more or less frequently than predicted by the neutral model (referred to as over-and underrepresented ASVs, respectively) and ASVs that followed the distribution of the neutral model (Figure 5d and e). For a small subset of ASVs a model fit could not be determined (from 3.7% in RS4 samples to 17.2% in RP1 samples, light green parts in Figure 5d), since those ASVs were below the detection limit or absent from S samples that were used as the seeding metacommunity in this analysis. Therefore, we assumed that these ASVs were also overrepresented in root-associated compartments compared to the bulk soil, although we kept them separate for additional analysis and refer to them as ‘overrepresented +’. While the model fit was similar for the RS4 and RS3 compartments, the migration rate *m* was lower for RS3. RS3 contains more ASVs with a larger positive deviation from the neutral model than RS4 (Figure 5d), meaning that their prevalence is higher than expected according to neutral processes. This indicates that while selection is equally important in RS4 and RS3, priority effects may play a bigger role in RS3 and there is less immigration than in RS4. All in all, microbial community composition near plant roots is primarily driven by selection, priority effects play an important role, and immigration from bulk soil is limited.

**Figure 5.**
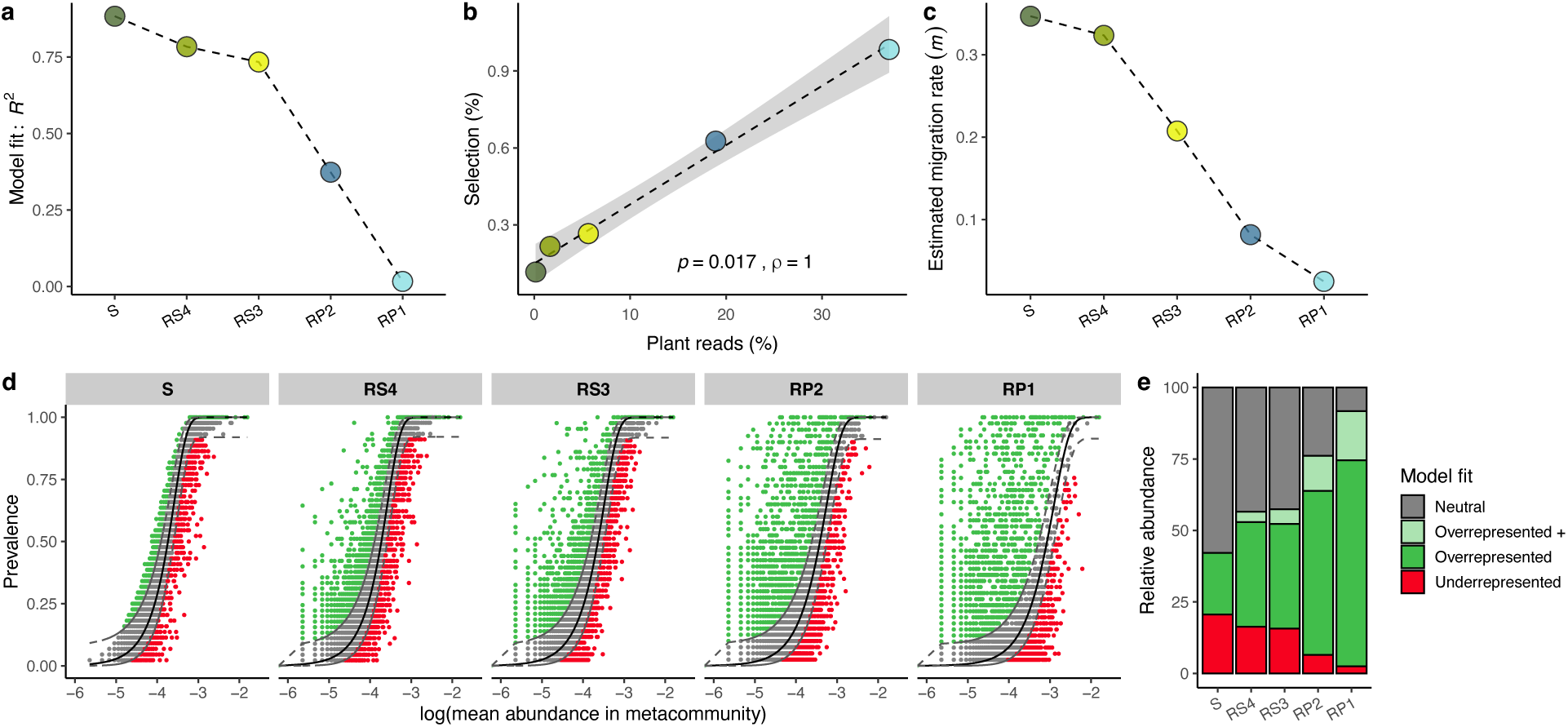
Selection-based community processes become more important closer to the root. **a** Neutral model fit (generalized *R*^2^) per compartment. A higher neutral model fit means less influence of selection-based community assembly processes. **b** Estimated migration rate per compartment. **c** The importance of selection, represented by the *R*^2^ of the model fit, as a function of the distance to the root, proxied by the number of plant reads per compartment (Spearman, *p* = 0.017, *rho* = 1). **d** Prevalence in compartment samples as a function of abundance in the bulk soil for each ASV per compartment, coloured by model fit. **e** The percentage of overrepresented ASVs (dark and light green bars) increases as distance to the root decreases, i.e., a larger part of the microbial community behaves non-neutral, suggesting a larger rhizosphere effect. Compartments are indicated by abbreviations: soil (S), rhizosphere-4 (RS4), rhizosphere-3 (RS3), rhizoplane-2 (RP2) and rhizoplane-1 (RP1).

### Phylogenetically similar microbes selected along the gradient

The proportion of overrepresented ASVs increased near the root, while that of the underrepresented ASVs, and ASVs that followed the neutral distribution decreased (Figure 5e). Neutrally distributed taxa are primarily generalists, widespread genera with opportunistic growth strategies, while taxa that are positively selected for are likely specialists adapted to a specific environment^49^. Therefore, we expected that the phylogenetic composition of the over- and underrepresented ASVs was different from the microbes whose abundances were described well by the model. To investigate this, we performed Non-metric Multidimensional Scaling (NMDS) analysis using the weighted UniFrac metric to visualize differences in community structure among the partitions of ASVs with different model fits per compartment (Figure 6). Pair-wise permutation analysis (999 permutations) revealed that all partitions had a distinct microbial community (PERMANOVA, *p* < 0.05). The phylogenetic composition of the partitions that diverged from neutral patterns remained relatively similar across the soil-root gradient, despite the changing composition of compartment communities as a whole. Among the overrepresented microbes are those genera that we identified previously that align well with the soil-root gradient, e.g. *Massilia* and *Niastella*. An interesting overrepresented+ genus is *Pseudomonas*, a well-known root coloniser^50–55^. Although *Pseudomonas* was not detected in bulk soil samples, several ASVs were found in RS3 and RP1 samples. Overall, taxa that get selected on the root are already visible in compartments further away from the root, albeit less abundant. Thus, although the rhizosphere effect is not yet clearly visible in the RS4 samples, a part of the microbes in this compartment is already influenced by the plant root and its exudates.

**Figure 6.**
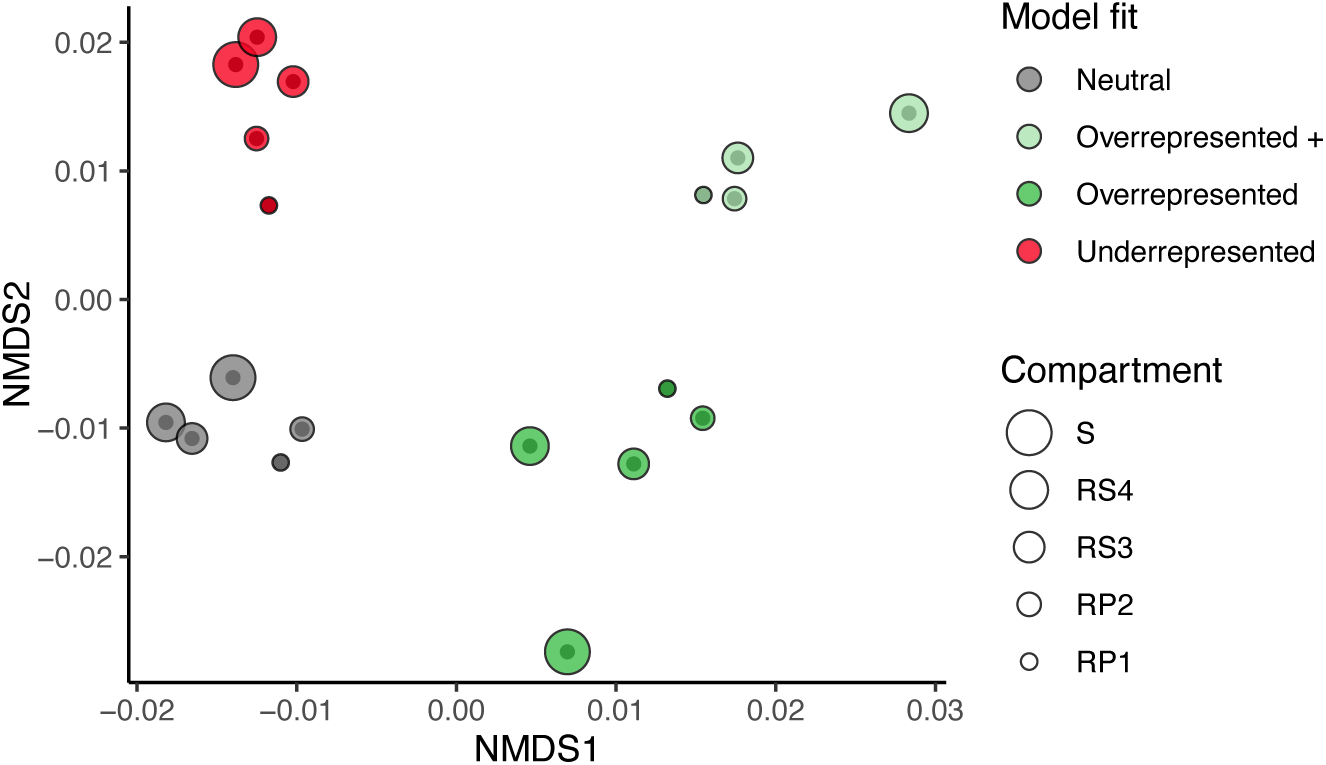
ASVs that are selected for or against along the soil-root gradient are phylogenetically distinct from ASVs that follow a neutral distribution. Partitions of overrepresented ASVs are shown in light and dark green, underrepresented ASVs in red and neutrally distributed ASVs in grey. Shape size indicates compartment, smaller symbols indicate less sampled soil. Compartment abbreviations: soil (S), rhizosphere-4 (RS4), rhizosphere-3 (RS3), rhizoplane-2 (RP2) and rhizoplane-1 (RP1).

### Rhizosphere microbes might use different growth strategies

Most microbes living in the proximity of plant roots are so-called ‘*r*-strategists’: copiotrophs that have high nutritional requirements and can exhibit high growth rates when resources are abundant^56^. Nutrient levels are higher in the rhizosphere compared to the bulk soil due to the secretion of compounds by plant roots. As a result, the concentration of nutrients is typically represented by a gradient, being the highest near the root and decreasing further away. Given our sampling strategy, we expected the average growth rate potential to go up gradually along the soil-root nutrient gradient, inversely related to root distance. We recently showed that rhizosphere-associated bacteria typically display higher growth rate potential and encoded more mobility mechanisms than soil bacteria, partly explaining their ability to reach and proliferate in the rhizosphere^37^. To assess whether growth rate potential can be modelled as a function of the distance to the root in addition to comparing the different biomes, we compared the median predicted minimal doubling time (PMDT) for each genus in our dataset (see Methods). For each compartment, we calculated the median PMDT as a weighted median of all PMDT values for all genera weighted by their relative abundances, and plotted them against the amount of plant reads as a proxy for root distance. We found that the growth rate potential of the microbiota was the highest in the two ‘rhizoplane’ compartments (RP2 and RP1) and ∼1.5 times higher than in S (Wilcoxon test, *p* < 0.01). The increase in growth rate potential in these rhizoplane compartments was effectively the largest, corroborated by the significant log-linear model fit (Linear model, log(y)∼x, *p* = 0.043, *R*^2^ = 0.791; Figure 7a). Growth rate potential flattens out towards the root, indicating that the maximum growth rate potential is reached. These results corroborate our earlier results that root-associated communities are enriched in microbes with a high growth rate potential compared to bulk soil^37^ and that the largest increase can be observed in the rhizosphere compartment.

**Figure 7.**
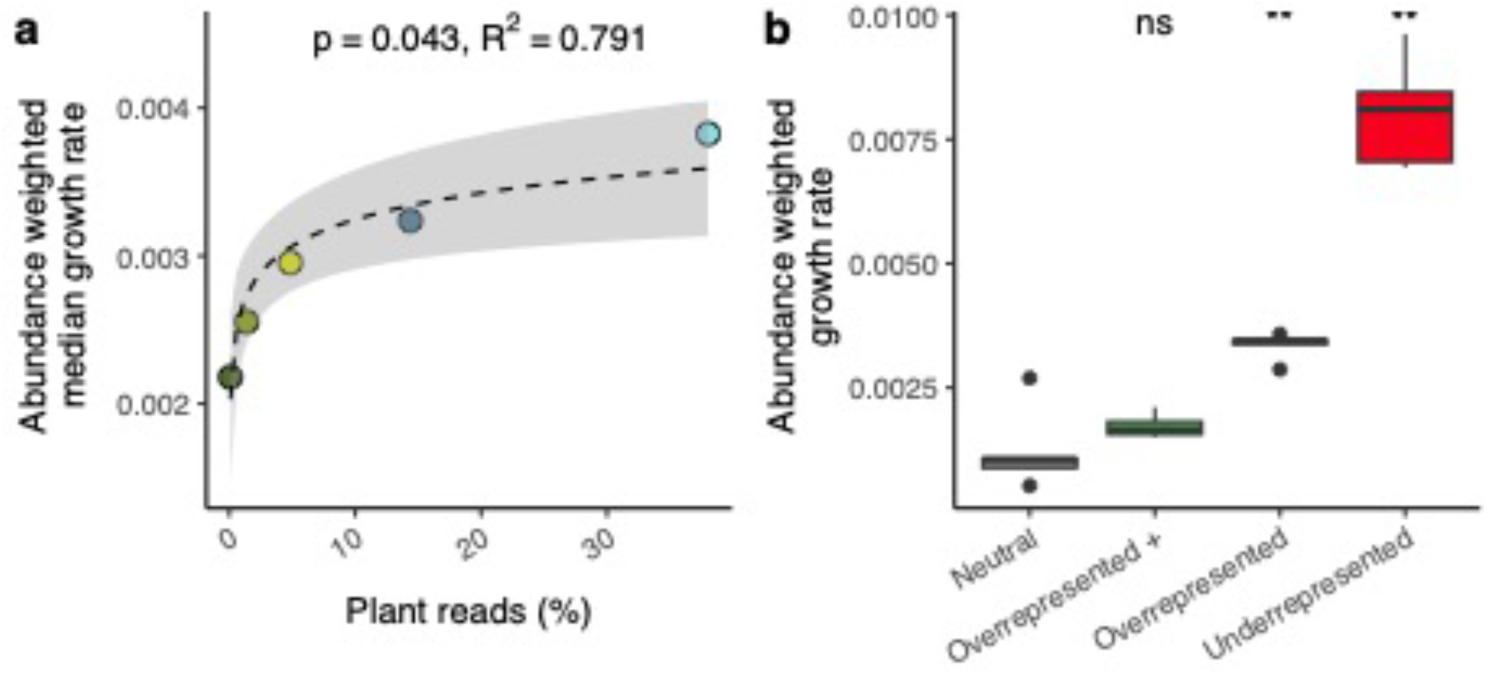
Growth rate potential increases closer to the roots, but rhizosphere-competent microbes have different growth strategies. **a** Abundance-weighted median growth rate potential per compartment is significantly correlated with the percentage of plant reads (Linear model, log(y)∼x, *p* = 0.043, *R*^2^ = 0.791). ‘Rhizoplane’ compartment microbes have on average a higher median growth rat potential as compared to bulk soil microbes (Wilcoxon, *p* < 0.01). **b** Abundance-weighted growth rate potential for microbes categorized by their neutral model fit. Compared to neutrally distributed microbes, microbes that are over- and underrepresented in the root-associated compartments display higher median growth rate potential (Wilcoxon test, *p* = 0.008).

Most likely, not only copiotrophs are able to survive in proximity of the root, but microbes can also use other strategies to colonize the rhizosphere and outcompete others. Therefore, we calculated the abundance-weighted median growth rate potential of taxa for the groups of ASVs based on their neutral model fits to assess whether over- and/or underrepresentation of microbes is linked to their respective growth rate potential. We found that the group of ASVs that were underrepresented in the root-associated compartments had the highest growth rate potential: ∼6.5 times higher than the neutrally distributed ASVs (Figure 7b; Wilcoxon test, *p* = 0.008). These microbes are less prevalent on the root than expected based on their abundance in soil, likely due to competition, mutual inhibition or by low tolerance to plant root-excreted antimicrobial compounds like coumarins^39^. Thus, being a potential fast grower is not sufficient to thrive in the rhizosphere, potentially due to competition with other fast growers. They might, however, still manage to colonize the root because they are generalist species that can survive in a wide range of environments^57^. In terms of their growth rate potential, the underrepresented ASVs were followed by ASVs that were overrepresented in the root-associated compartments with average growth rate potential ∼2.5 times higher than neutrally distributed ones (Wilcoxon test, *p* = 0.008). Overrepresented ASVs make up the largest proportion of bacteria in the ‘rhizoplane’ compartments, and likely complement their relatively high growth rate potential with other traits that make them highly rhizosphere competent. Interestingly, genera within the overrepresented+ class had similar growth rate potential as the ASVs whose distribution followed the neutral model (Wilcoxon test, *p* > 0.05). While these ASVs were not detected in the bulk soil and are predicted to be relatively slow growers, they are still able to colonize the root in high numbers (**Error! Reference source not found.**). These microbes, amongst them the previously noted genus *Massilia*, are probably specialist species that have specific traits not related to their growth rate potential, like competitive ability, that help them survive and thrive in the rhizosphere.

## Discussion

In this study, we employed a reproducible sampling approach to harvest the roots and adhering soil of individual Arabidopsis plants cultivated in different soil batches. Using increasingly stringent cleaning and washing steps, we sampled the soil-to-root continuum in five consecutive compartments, including the unplanted bulk soil itself. This enabled us to investigate microbial community composition and community assembly processes along this continuum with high resolution, gaining critical insights in the microbial dynamics in this hotspot of microbial activity. As we move closer to the root, we observed that bacteria from the *Proteobacteria* and *Bacteroidetes* replaced other phyla and that microbes that were below the detection limit in soil can be the most abundant species in the rhizosphere or rhizoplane. Despite different initial soil microbiomes, rhizosphere and rhizoplane microbiomes converge when we considered microbial phylogenetic relatedness, supporting earlier work showing that specific bacterial families and genera are repeatedly associated with plant roots^19^. It also shows that plant roots impact their local environment in a robust and reproducible manner, exerting selective pressure on the surrounding microbes likely to their own benefit. Nevertheless, we also observed that, in addition to selection pressure- or niche-based community assembly, also priority effects and alternative growth strategies can contribute to shaping the rhizosphere microbiome.

### Comparative analysis of the rhizosphere microbiome across plants, soils and samples requires reproducible sampling

Our findings highlight the importance of careful consideration when sampling roots in rhizosphere studies. Different sampling methods can lead to varying conclusions regarding the strength of the rhizosphere effect and its root-associated microbiome and assembly processes. In the current study, α-diversity was significantly lower in the ‘rhizoplane’ compartments when compared to bulk soil. We expect that the reduction in α-diversity is accompanied by a reduction in functional gene richness, as taxonomic and functional diversity are inherently linked^58^. The same effect was seen for β-diversity, as the community composition diverged from bulk soil while closing in on the root. More importantly, both diversity parameters correlated significantly with the distance to the root – approximated in our study by the number of plant-derived reads – emphasizing the need to treat the rhizosphere as an environmental continuum or gradient. Earlier cultivation-dependent and -independent studies recognized lower α- and higher β-diversity in the rhizosphere compared to bulk soil, but to what extent depended heavily on the sampling method. In studies on Arabidopsis, analysis of species richness – a proxy for α-diversity – in the rhizosphere compartment displayed a slight decrease^59^ or no change at all^19,22,60,61^. More often, α-diversity decreased significantly only when examining the rhizoplane or the endophytic compartment^18,19,60,61^, which compares best to our ‘rhizoplane’ compartments RP2 and RP1^71^. In an earlier study on the Arabidopsis rhizosphere effect, rhizosphere and endophytic communities were found both to be different from bulk soil, and this effect was largest for the endophytic community^19^. The compartment effect found in Lundberg et al. (2012) (rhizosphere vs. endosphere) aligned well with the largest principal component of the PCoA. Comparable to the variation between our different experiments, soil batch was represented by the second principal component^19^. These findings are comparable to results from Bulgarelli and co-authors (2012). Here, analysis of sample-to-sample β-diversity revealed that all root samples were distinct from rhizosphere and bulk soil samples, irrespective of soil type. For rhizosphere and soil samples this distinction was less clear^18^. Two other studies that compared the rhizosphere and endosphere show similar trends: the difference between rhizosphere and soil microbiomes (based on the PERMANOVA-test *R*^2^) was relatively small, while for the endosphere-soil comparisons *R*^2^-values were larger, indicating more dissimilar communities^22,61^. Our rhizoplane-to-soil β-diversity compartment effect sizes are smaller than those reported by Schneijderberg *et al*.^22^, likely due to the spread between samples of different experiments. Experiment-specific *R*^2^ values ranged from 0.42 – 0.72 for RP1 samples, which are comparable to theirs.

### Rhizosphere-competent *Massilia* species are rare in soil, yet abundant on plant roots and understudied

The gradual change in community composition along the soil-root gradient was mostly caused by a doubling in abundance of *Proteobacteria* closer to the roots. Members of this phylum include fast-growing, generalist *r*-strategists that do well in environments where organic resources are abundant^56^. They are often found enriched on the roots of Arabidopsis^18,19,21,22,62^ and many other plants including crops like maize^63,64^ and wheat^65–67^. Within the *Proteobacteria*, we found members of the *Betaproteobacteria* to increase in particular, and within this class, the family *Oxalobacteraceae* was most prominent. *Oxalobacteraceae* are good rhizosphere and even better endosphere colonizers^18,60^. Of these, the genus *Massilia* correlated most strongly with the first axis in our PCoA which distinguished samples of different compartments. It was nearly absent in the bulk soil, but overrepresented closer to the roots where it occurred in high abundance. *Massilia* species are generally good colonizers of biological surfaces, and often found enriched in the rhizosphere^21,52,62,67^. They associate mostly with older parts of the plant root^68^, suggesting that they do not necessarily need highly metabolically active cells for survival. This might give them a competitive advantage over other microbes that need to be in proximity to root tips to grow. Our fine-grained sampling focused on complete roots of 4-week-old Arabidopsis plants, and we are thus unable to differentiate microbial communities from young or older parts of the roots. However, *Massilia* species have been reported to display typical plant-growth-promoting capabilities^68^ in association with older root tissues. For example, in nitrogen-poor soil, *Massilia* isolates were able to promote maize shoot growth and nitrogen accumulation with induction of lateral root formation in lateral rootless mutants^69^, a process that takes place in the older parts of the root. A specific *Massilia* ASV was found to be associated with pathogen-infected Arabidopsis plants^70^, and it was also identified as a ‘hub’ species in the wheat rhizosphere, suggesting it plays an essential role in the assembly and functionality of plant-associated communities^67^. *Oxalobacteraceae*, and specifically the genus *Massilia*, are extremely rhizosphere competent and might fulfil essential functions within the rhizosphere microbiome. To date only 139 *Massilia* spp. genome sequences are deposited in the NCBI GenBank database. Compared with well-known, and well-studied rhizosphere-competent genera like *Bacillus* (9,322 genomes), *Pseudomonas* (8,000 genomes excluding the opportunistic human pathogen *Pseudomonas aeruginosa*), *Rhizobium* (1,500 genomes), and *Agrobacterium* (697 genomes) this highlights our limited understanding and characterization of this genus. Such is further corroborated by the much smaller number of scientific studies that can be found via NCBI PubMed and that evaluated the interaction between members of the *Massilia* genus and plants (1,198 studies) as compared to that involving abovementioned genera (*Pseudomonas*: 106,543; *Bacillus*: 100,729; *Agrobacterium*: 51,095; *Rhizobium*: 20,302 studies).

### Deterministic processes drive root microbial community assembly

Our research points towards niche-based community assembly processes driving the rhizosphere effect, resulting in phylogenetically similar microbial communities on the root. These consist for an important part of organisms from fast-growing generalist taxa^37^. Few studies try to quantify microbial community assembly in the root environment, and results are variable. One study on the soybean rhizosphere suggested that bulk soil microbiomes were governed solely by neutral processes, whereas rhizosphere samples showed deterministic dynamics^71^. Another study, also on soybean rhizospheres, found that deterministic processes were more important in bulk soil than in the rhizosphere and endosphere^30^. A third soybean study found that neutral processes shaped rhizo-and endosphere microbiomes, although those were not compared to bulk soil samples^72^. In long-term cultivated wheat, Fan and co-authors (2017) found that deterministic assembly processes dominated bacterial community composition in plant and soil compartments^29^. All these studies used different methods to quantify the importance of niche and neutral community assembly processes, including differences in sampling strategy and analysis, so a one-to-one comparison is complicated. We expect that extending our approach of incrementally sampling the rhizosphere to other plant species, will help elucidate not only what assembly processes dominate the rhizosphere, but also what microbes are selected for or against.

### Priority effects drive compartment differentiation across plants and experiments

When we conducted PCoA using various β-diversity metrics, samples clustered by noticeably different patterns. Samples from the three different experiments diverged in compartments closer to the root when we used Bray-Curtis, but not when we used weighted UniFrac. This means the number of different ASVs increased, while their phylogenetic diversity remained stable from soil to root. Additionally, the neutral model predicted that migration rates decreased near the roots, indicating that there is enhanced growth of the local community and less migration from the bulk soil. This is consistent with the higher growth rate potential of rhizosphere-associated bacteria^37^ and highlights the importance of priority effects in the colonization of the rhizosphere^74^. Using synthetic communities and sequential inoculations, Wippel and co-authors (2021) showed that the order of arrival was indeed a major factor determining the final microbial community on *Arabidopsis* and *Lotus japonicus* roots^61^. The effect of secondary inoculations on the community depended on the context, e.g., the origin of the inoculated community, the plant compartment, and the nutritional status of the plant, which might all influence the competitive ability of present microbes. Thus, understanding the effects of timing on rhizosphere community assembly will be crucial for gaining a complete picture of these complex systems.

In conclusion, our study investigated the rhizosphere effect in Arabidopsis along a soil-root gradient, utilizing three distinct experiments with comparable sampling methods. Microbial communities displayed a gradual shift in composition along the gradient, with *Proteobacteria* and *Bacteroidetes* dominating the rhizoplane compartments. The rhizosphere effect, indicated by decreasing α-diversity and distinct β-diversity patterns, strengthened towards the root. The enrichment of specific taxa on roots, such as *Massilia* and *Devosia*, revealed phylogenetic clustering and suggested a non-random selection of microbes. Priority effects became more pronounced closer to the root, influencing community structure. Modelling using the Sloan Neutral Community Model indicated that the root-associated communities were primarily shaped by non-neutral processes like selection, with priority effects playing a larger role in compartments with limited migration from bulk soil. Additionally, growth rate potential analysis suggested that taxa that were underrepresented on the root had the highest growth rate potential, emphasizing the complex interplay of (growth) strategies in the rhizosphere. Our discoveries advance our comprehension of the intricate dynamics involved in rhizosphere microbiome assembly, underscoring the critical role of sampling strategies. This insight is particularly pertinent for future research delving into plant-microbe interactions, highlighting the significance of methodological approaches in unraveling the complexities of the rhizosphere microbiome.

## Acknowledgements

The authors would like to express their gratitude to the members of the Utrecht University Plant-Microbe Interactions lab and Metagenomics Group (MGX) for their valuable discussions. We acknowledge the Utrecht Sequencing Facility (USEQ) for providing sequencing service and data. USEQ is subsidized by the University Medical Center Utrecht and The Netherlands X-omics Initiative (NWO project 184.034.019). This study was supported by the NWO Green II Grant no. ALWGR.2017.002 (S.W.M.P., J.J.S.G., R.d.J.) and the Novo Nordisk Foundation Grant no. NNF19SA0059362 (R.d.J.). B.E.D. was supported by the European Research Council (ERC) Consolidator grant 865694: DiversiPHI, the Deutsche Forschungsgemeinschaft (DFG, German Research Foundation) under Germany’s Excellence Strategy – EXC 2051 – Project-ID 390713860, and the Alexander von Humboldt Foundation in the context of an Alexander von Humboldt-Professorship funded by the German Federal Ministry of Education and Research.

## Author contributions

**Sanne W.M. Poppeliers:** Conceptualization, Methodology, Analysis, Visualization, Writing – Original Draft. **Juan J. Sánchez-Gil:** Investigation, Writing: Review and Editing. **José L. López:** Analysis of growth rate potential data. **Bas E. Dutilh:** Supervision, Funding acquisition, Writing: Review and Editing. **Corné M.J. Pieterse:** Supervision, Funding acquisition, Writing: Review and Editing. **Ronnie de Jonge:** Supervision, Project administration, Funding acquisition, Writing – Review and Editing.

## Supplementary material

**Supplementary Figure 1.**
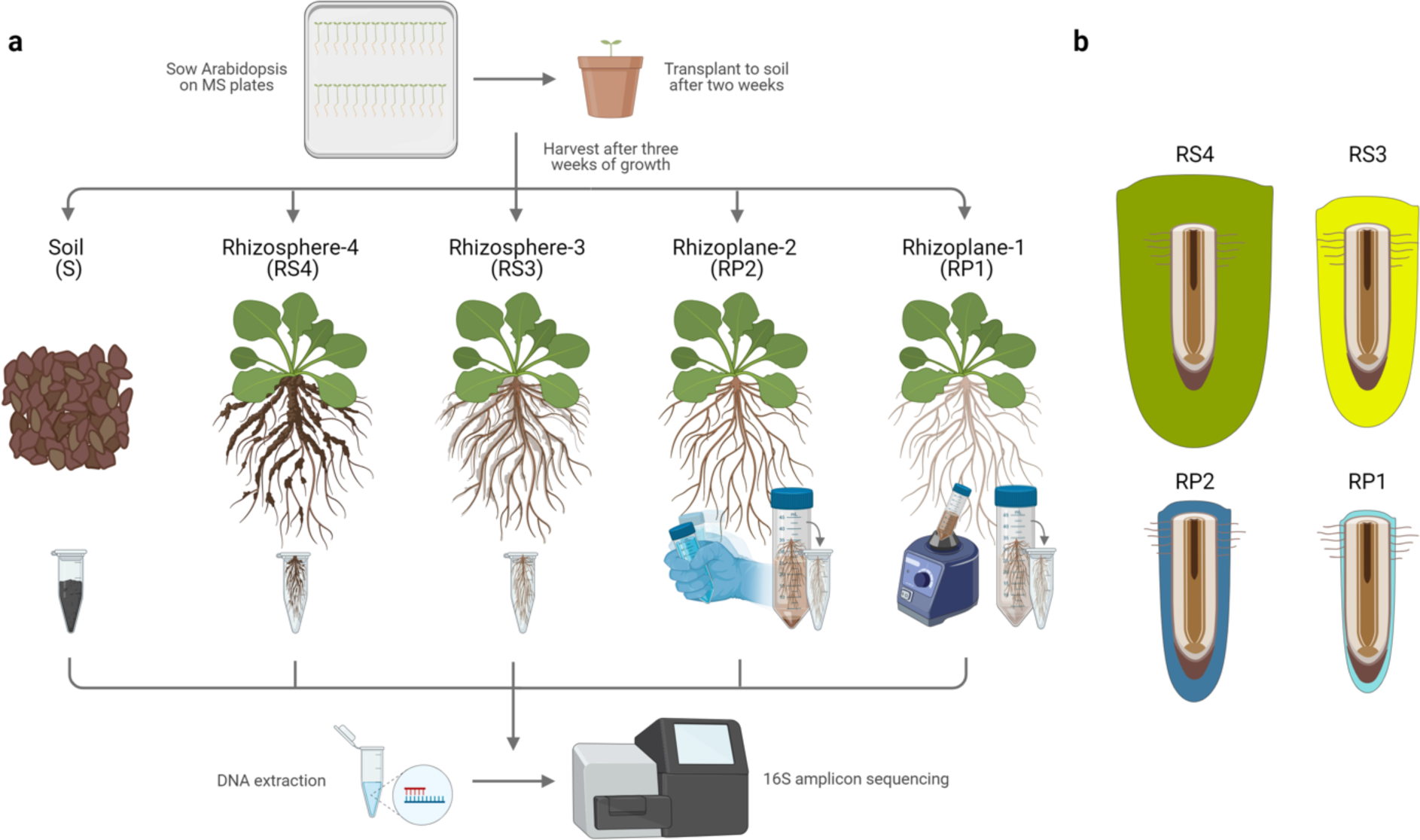
Schematic overview of the experimental setup of the rhizosphere effect experiments. **a** Surface-sterilized Arabidopsis seeds were sown on plates with Murashige and Skoog medium. After two weeks seedlings were transplanted to soil and grown for another three weeks until harvest. Unplanted samples were considered bulk soil (S). About 0.25g of soil was taken per pot. Then, for all root compartments, roots were harvested in different ways and collected in 2-mL Eppendorf tubes. For rhizosphere-4 samples (RS4), roots were taken out of the soil and gently shaken until most loose soil fell off. This compartment contained the widest zone of soil environment around the roots and its microbial community is likely to show the closest resemblance to that of the bulk soil. For rhizosphere-3 (RS3), roots were harvested similarly, but additionally they were cleared of most loose soil by tapping them on paper and stripping them from soil as much as possible. The rhizoplane-2 (RP2) roots were harvested as RS3, and additionally gently shaken in a 10 mM MgSO4 solution. For rhizoplane-1 (RP1), roots were additionally vortexed twice in a phosphate-Silwet buffer. As a result, in each consecutive compartment from RS4 to RP1 we sampled bacterial communities more closely associated with the root, or not influenced by the root at all (S). We consider that RS4 and RS3 may be compared to what other studies often refer to as ‘rhizosphere’, while RP2 and RP1 could be considered ‘rhizoplane’. Notably, our sampling strategy also included the roots themselves, sometimes referred to as ‘endosphere’. **b** Schematic view of how the sampling strategy results in a gradual decrease in soil and microbes attached to the roots.

**Supplementary Figure 2.**
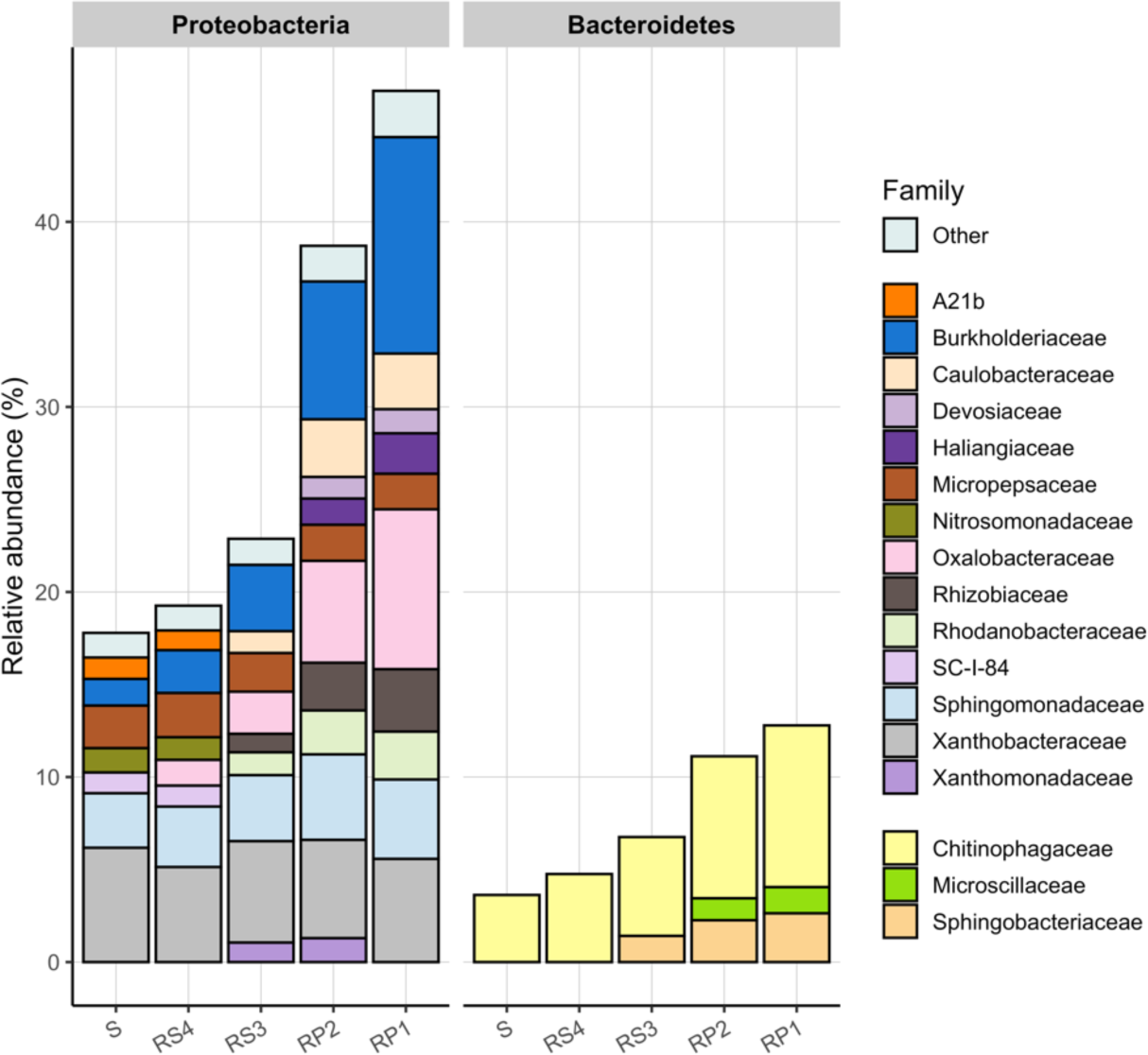
*Burkholderiaceae* and *Oxalobacteraceae* (Proteobacteria), and *Sphingobacteriaceae* and *Chitinophagaceae* (Bacteroidetes) increase in abundance closer to the root. The relative abundance of families within the Proteobacteria and Bacteroidetes that have an abundance of more than 1%, colored by family, shown per compartment. Compartments are indicated by abbreviations: soil (S), rhizosphere-4 (RS4), rhizosphere-3 (RS3), rhizoplane-2 (RP2) and rhizoplane-1 (RP1). ‘Other’ taxa are not classified on the family level.

**Supplementary Figure 3.**
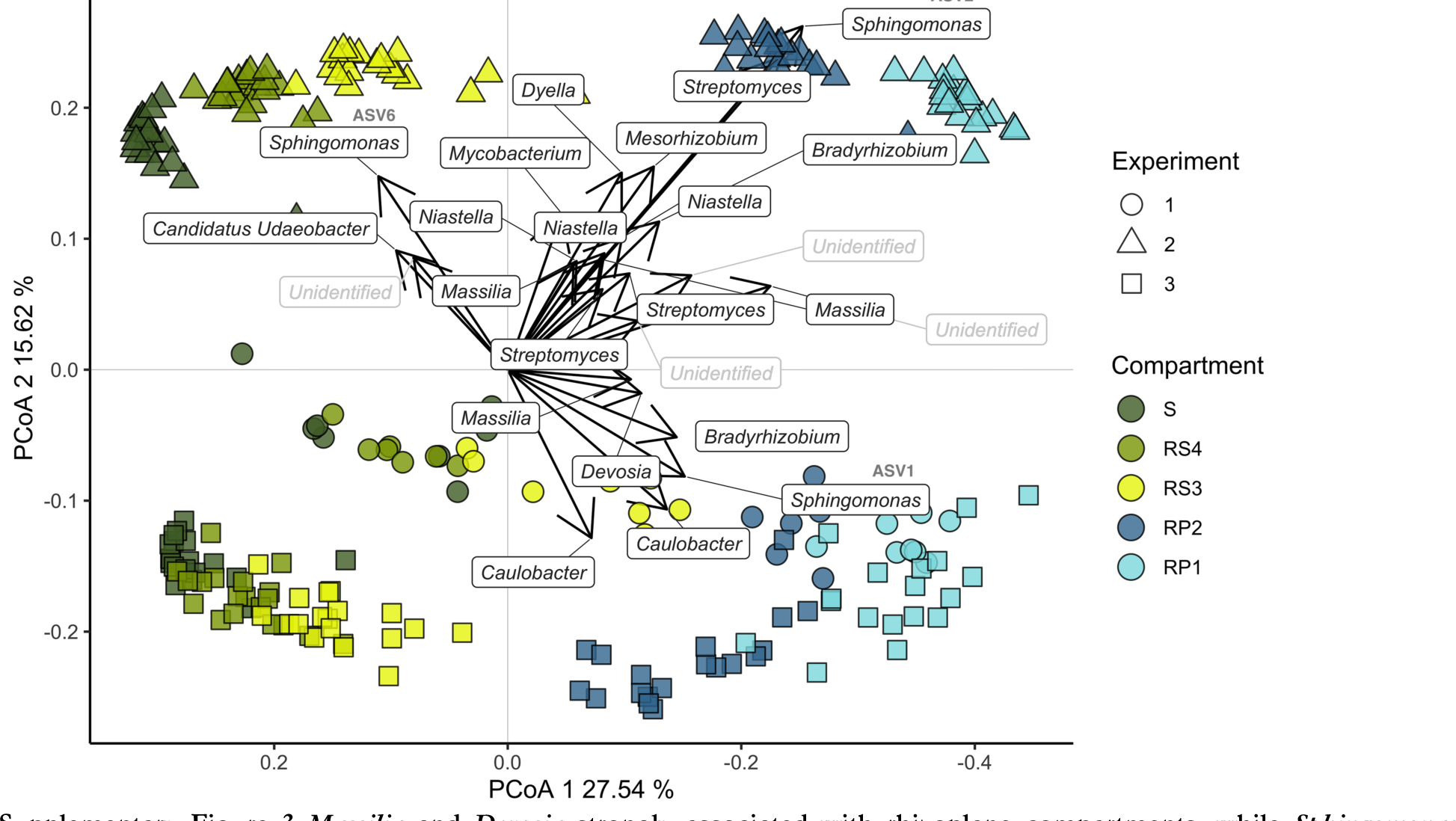
*Massilia* and *Devosia* strongly associated with rhizoplane compartments, while *Sphingomonas* contributes most to the ordination. PCoA biplot using the Bray-Curtis distance metric where arrows show the 25 ASVs that contribute most to the ordination. The longer the arrow the larger the contribution. Arrows that align best with PCoA1 (grey line) are correlated strongest with the soil-root gradient. Compartments are indicated by abbreviations: soil (S), rhizosphere-4 (RS4), rhizosphere-3 (RS3), rhizoplane-2 (RP2) and rhizoplane-1 (RP1). ‘Unidentified’ taxa are not classified on the genus level.

**Supplementary Figure 4.**
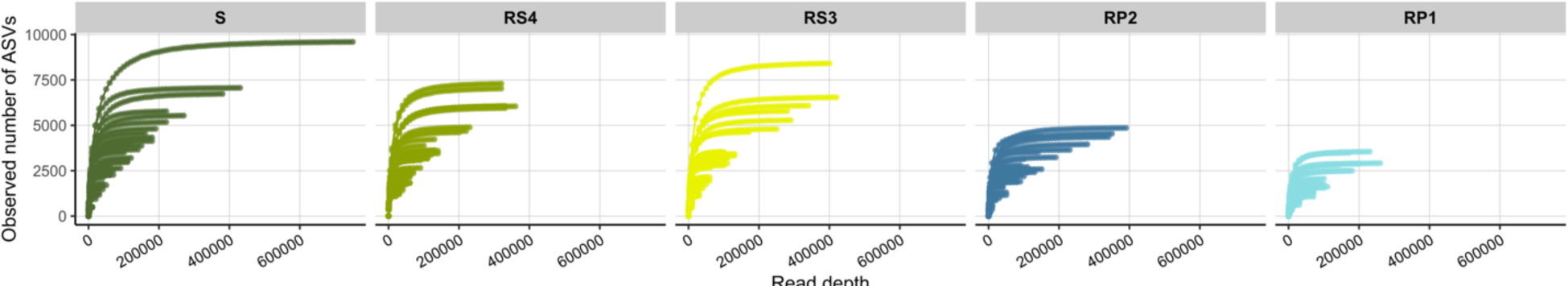
Saturation is nearly reached in almost all samples. Rarefaction curve for all samples, coloured by compartment.

**Supplementary Table 1.**
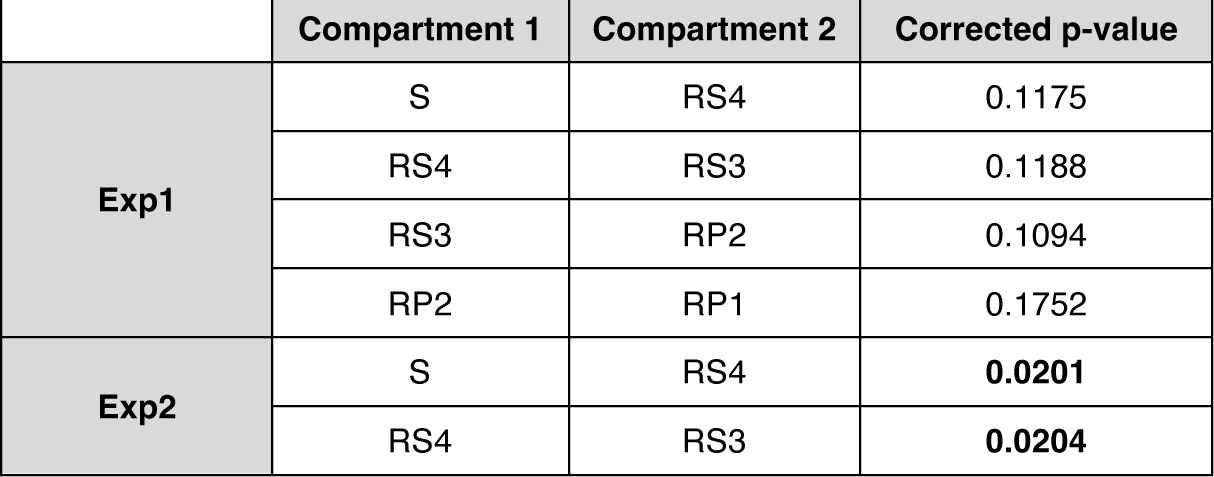
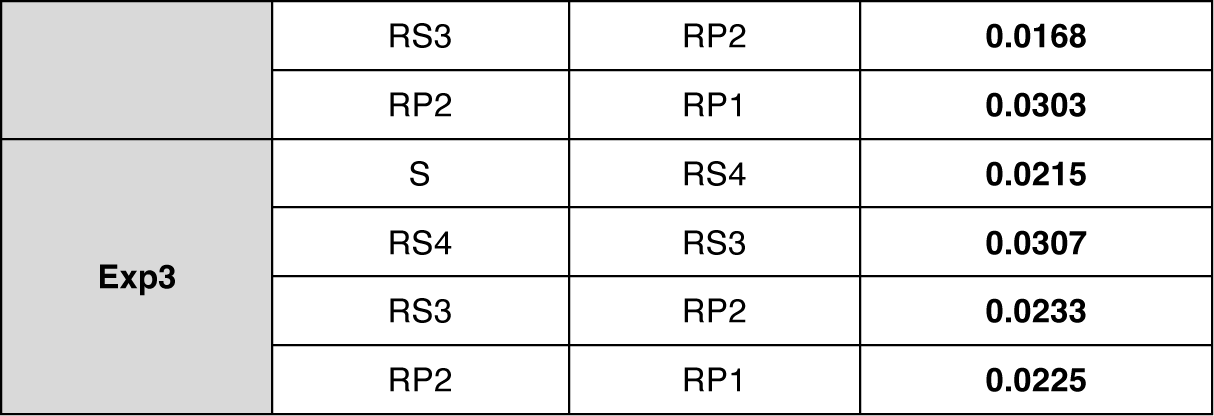
Amount of plant reads increases towards the roots. Dunn’s post-hoc test results after Kruskal-Wallis test comparing the amount of plant reads between neighbouring compartments. For Exp1, differences between neighbouring compartments are smaller, therefore not significant (Figure 2b).

**Supplementary Table 2.**
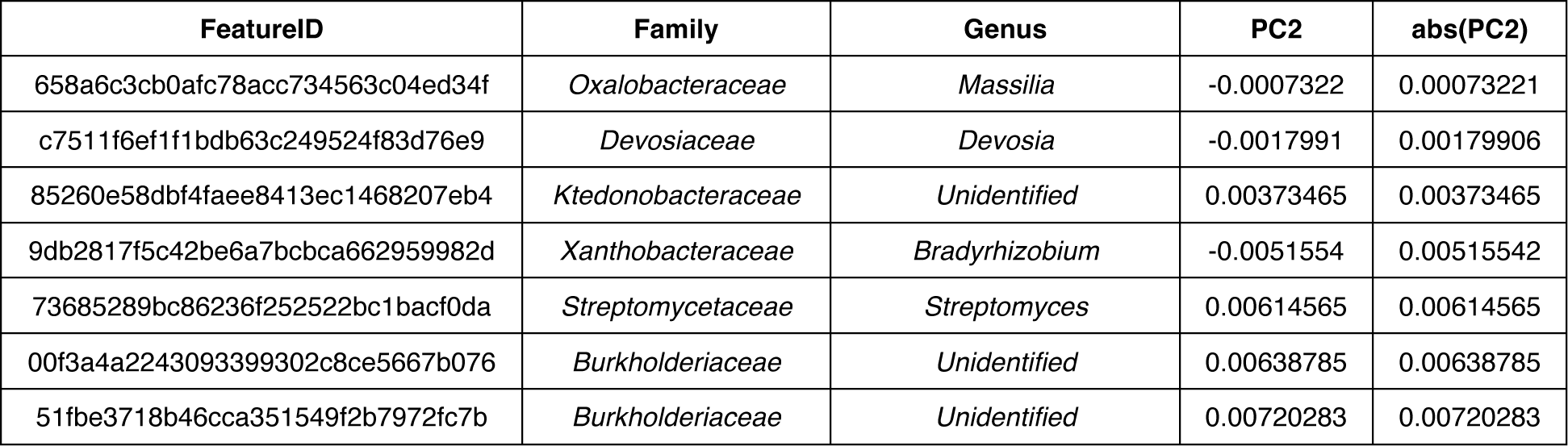
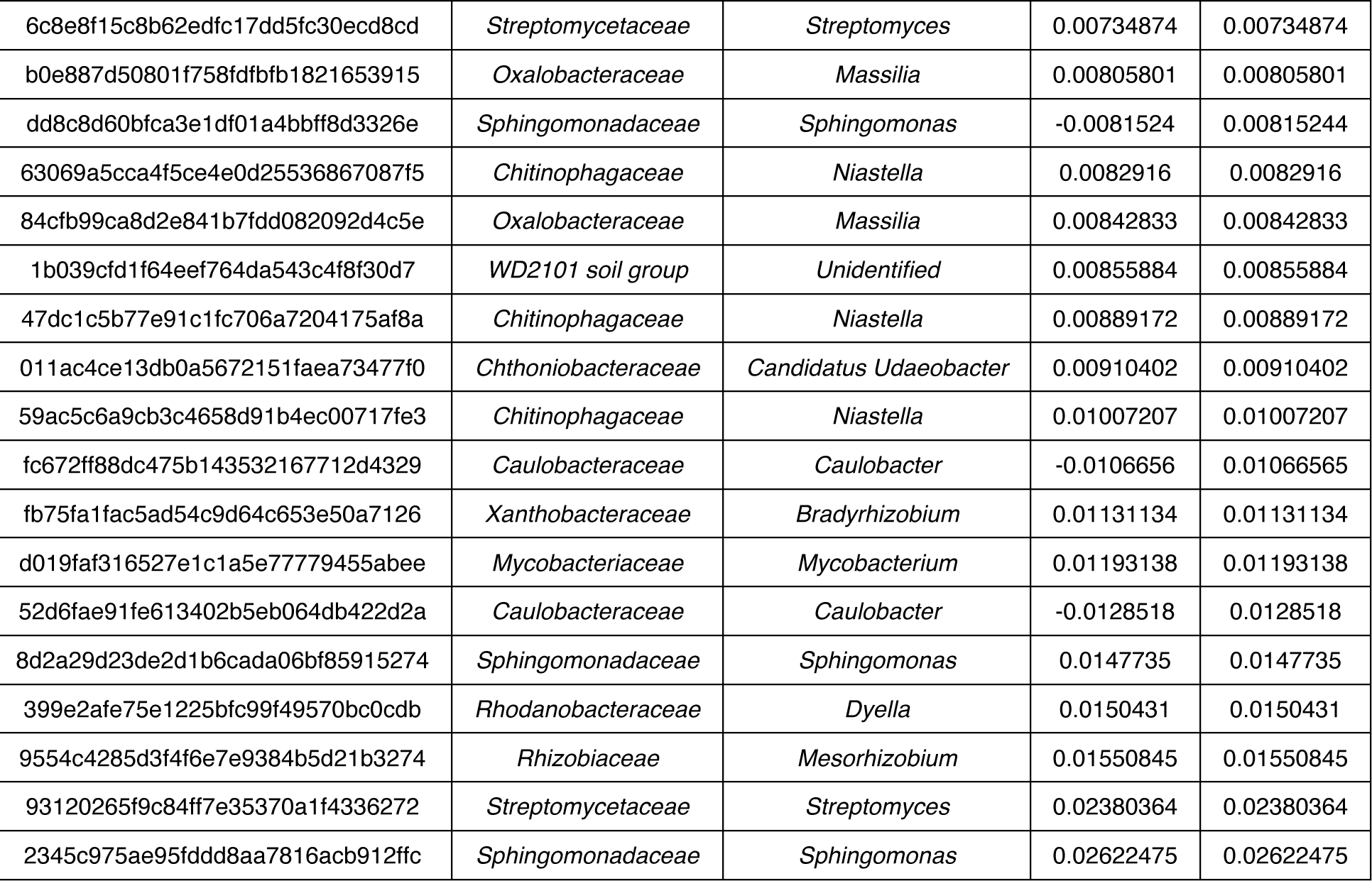
*Massilia* and *Devosia* ASVs most strongly associated with the rhizosphere effect. Absolute PC2-values indicate how strong an ASV correlates with PC1. The table is ranked from highest to lower correlation.

**Supplementary Table 3.**
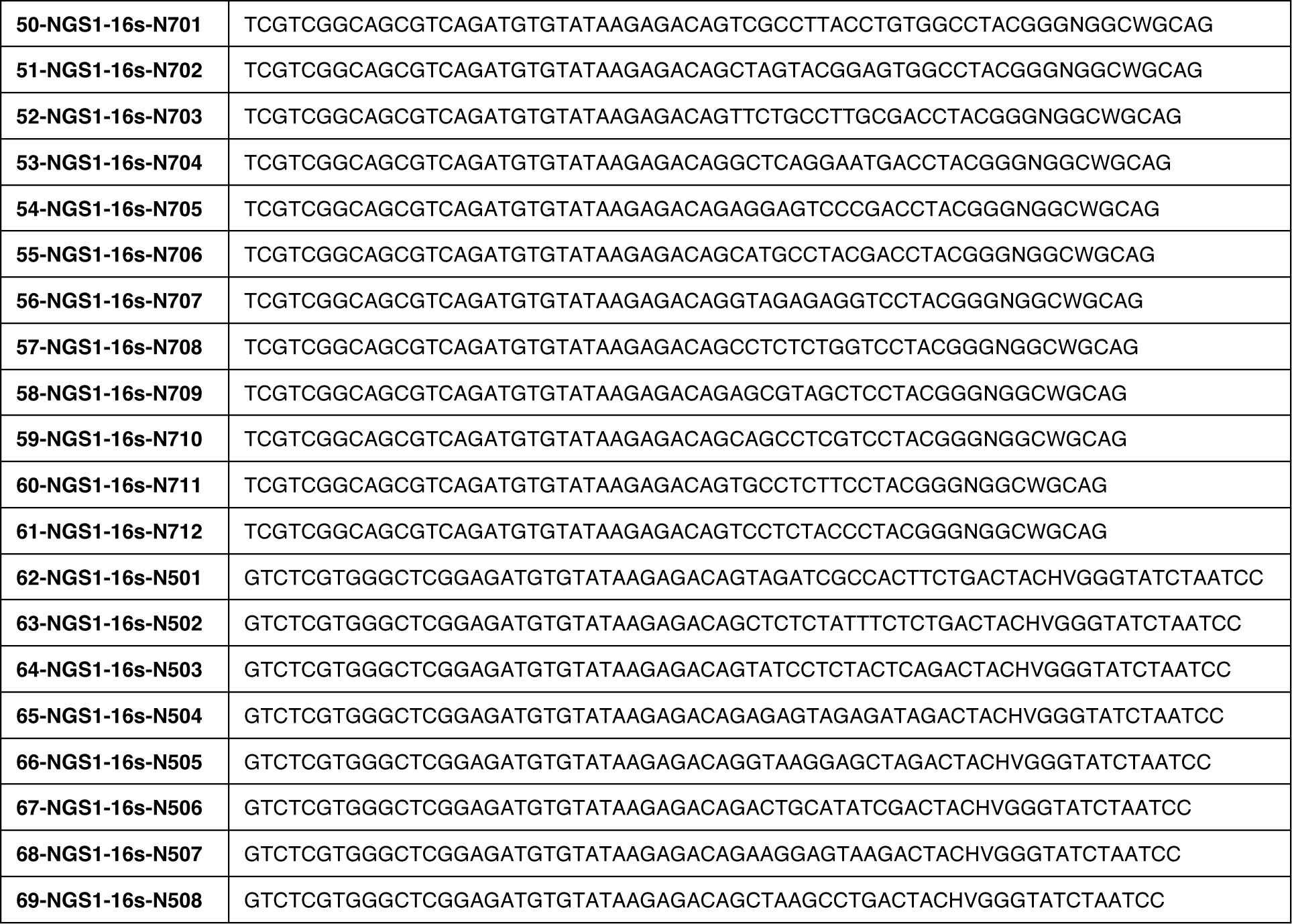
Phasing primers used for amplification of the hypervariable V3-V4 region of the 16S rRNA gene.

